# Abiotic cues modulate exopolysaccharides to influence type VI secretion system-mediated antagonism

**DOI:** 10.64898/2026.06.18.733139

**Authors:** Yi-Chieh Wang, Li-Kang Sung, Chih-Feng Wu, Jeff H. Chang, See-Yeun Ting, Erh-Min Lai

## Abstract

The type VI secretion system (T6SS) enables Gram-negative bacteria to inject toxic effectors into neighboring cells, mediating contact-dependent antagonism and interbacterial competition. How the T6SS-mediated attack responds to environmental cues varies and remains unclear among different bacteria. Here, using *Agrobacterium fabrum* C58, a soil-borne phytopathogenic bacterium, we investigated the impact of osmolarity, moisture, surface stiffness, and glucose on T6SS-mediated antagonism. We show that these abiotic factors influenced the production of two polysaccharides, cyclic-β-(1,2)-glucan (CβG) and succinoglycan (SG), and modulated the T6SS-killing outcome. Mechanistically, high osmolarity inhibits CβG production, thereby enhancing the expression and secretion of the T6SS. In contrast, SG biosynthesis, in response to moisture, surface stiffness, and glucose, does not impact T6SS expression and function but decreases T6SS-mediated killing efficacy. Electron microscopy revealed that SG creates a physical barrier between bacterial cells. Such physical distancing not only hinders the T6SS attack from *Agrobacterium*, but also confers protection against other competitors at both intra- and inter-species levels. Our results unravel the complexity of how specific environmental factors modulate the contact-dependent antagonism and highlight a balance between offensive and defensive behaviors.

**Significance Statement:** Bacteria live in polymicrobial communities where they often need to fight off competitors to survive. One well-characterized weapon is the type VI secretion system (T6SS), a nanomachine mediating contact-dependent antagonism. In this study, we aim to study how the T6SS attack is influenced by environmental cues. We discovered that carbon sources, osmolarity, and surface stiffness modulate the T6SS efficacy through the secretion of sugar chains outside the cell. This sugar secretion increases the physical distance between cells, protecting against T6SS-mediated attack. However, such physical distancing also hinders the efficacy of the T6SS attack originating from *Agrobacterium* itself. Our results reveal a previously understudied offense-defense tradeoff between EPS production and T6SS-mediated attack. This finding underscores the intricate balance between bacterial offense and defense strategies.

## INTRODUCTION

Bacteria commonly coexist in complex communities in the environment, where intense competition occurs (1). To compete for nutrients and ecological niches, bacteria adopt various defense or offense strategies. The type VI secretion system (T6SS) is a well-characterized nanomachine that mediates contact-dependent antagonism (2). It is widespread in Gram-negative bacteria, including many environmental strains as well as animal and plant pathogens (3). T6SS-dependent antagonism is intricately regulated at transcriptional, translational, and post-translational levels by abiotic and biotic cues (4). For example, in the squid light organ, where the symbiont *Vibrio fischeri* frequently encounters competitors, the viscous environments directly regulate expression of its T6SS (5). A mildly acidic to neutral pH further promotes cell-to-cell contact, which is required for contact-dependent killing (5). Variations of T6SS regulation are also observed across different species or even among the same species (6, 7). In clinically isolated *V. cholerae*, the T6SS is involved in successful gut colonization but remains inactive in standard lab conditions. In contrast, environmental isolates often exhibit constitutive T6SS-dependent killing. These exemplify how bacteria fine-tune T6SS regulation in response to distinct ecological contexts.

T6SS is a multipotent weapon capable of delivering various effector proteins that target a broad range of cellular components, including cell envelopes, energy sources, and nucleic acids (8). To prevent intoxication from self or neighboring sister cells, the attackers produce the cognate immunity proteins. Previous studies have revealed that in environments where the competition is intense, bacteria may harbor orphan immunity gene arrays to protect themselves against external attacks from competitors (9, 10). However, it is difficult to acquire all cognate immunity genes due to the diversity of T6SS toxins, and bacteria have adopted different strategies for protection against external attacks from competitors.

One known immunity protein-independent protection is exopolysaccharides (EPS), forming a physical barrier provides an effective means of defense against these attacks (11–13). In *Salmonella enterica*, for instance, lipopolysaccharide (LPS) acts as a barrier to T6SS-mediated attack (14). In *Vibrio cholerae*, production of capsular polysaccharide (CPS) reduces susceptibility to T6SS-mediated killing, while self-inflicted T6SS attacks remain unaffected (11). This suggests that, rather than merely separating bacteria from competitors, the capsule may act as armor that protects cells from external attacks. Capsule formation has also been identified as a widespread defense strategy in *Enterobacter* and *Klebsiella*, and such traits can be acquired through horizontal gene transfer (13). However, another study on *Acinetobacter baumannii* reported that the capsule formation reduces both the efficacy of the T6SS attack from itself or its competitors (15). This highlights the diversity of impacts that the CPS has on T6SS-mediated interactions, where the architecture, biophysical, and biochemical properties of capsules may determine the outcome in contact-dependent antagonism. In addition to the surface-bound LPS and CPS, secreted EPS also contribute to protection (12). These EPS form a slime layer surrounding the cells and facilitate biofilm formation. Notably, EPS production provides collective protection against T6SS attack, not only to producers but also to nearby non-producers (12). This suggests that EPS serves as a public good that modulates microbial interaction. However, it remains unclear whether secreted EPS influences T6SS activity originating from the same cell.

Agrobacteria are a group of bacteria with members capable of infecting plants to cause crown gall or hairy root diseases (16, 17). T6SS is present in a subset of agrobacteria and known to function as an antibacterial weapon for competitive advantages *in planta* and *in vitro* (7, 18–20). Using *Agrobacterium fabrum* C58 as a model, previous studies demonstrated that T6SS is upregulated in acidic conditions via ExoR-ChvG/ChvI two-component system (21, 22). This acidic upregulation may occur in the rhizosphere, where *Agrobacterium* encounters competitors and frequent attacks. Similar to T6SS, production of succinoglycan (SG), a predominant secreted EPS contributing to biofilm formation, is increased under acidic conditions through the ExoR-ChvG/ChvI pathway (23). This highlights the possibility of interplay between EPS production and T6SS-mediated competition, and leads us to hypothesize that *Agrobacterium* coordinates various physiological processes, including offense and defense responses, to thrive in the environment.

In this study, we investigate how abiotic cues and EPS production influence T6SS expression and the competition outcome of *Agrobacterium*. We found that the production of cyclic-β-(1,2)-glucan (CβG) negatively regulates T6SS activity, and demonstrated that SG offers physical distancing that compromises the T6SS efficacy. Our findings highlight an offense-defense tradeoff involving EPS production and contact-dependent antagonism.

## RESULTS

### Hyper-osmolarity and deficiency of β-(1,2)-glucan promotes T6SS antibacterial activity

To investigate whether and which environmental cues regulate agrobacterial T6SS, we used *Agrobacterium fabrum* C58 as a model and performed interbacterial killing assays under various conditions (Fig.1). We first tested whether *Agrobacterium* T6SS is regulated by osmolarity, and performed a killing assay against *E. coli* on LB agar plates supplemented with different concentrations of NaCl. We found that *E. coli* prey survival against *A. fabrum* C58 wild type (WT) decreased as NaCl concentrations increased (Fig.1A), while prey survival against the Δ*tssL* mutant, a T6SS-deficient strain, remained the same across concentrations. This suggests that the T6SS-killing outcome is enhanced by higher osmolarity or salinity.

**Fig 1.**
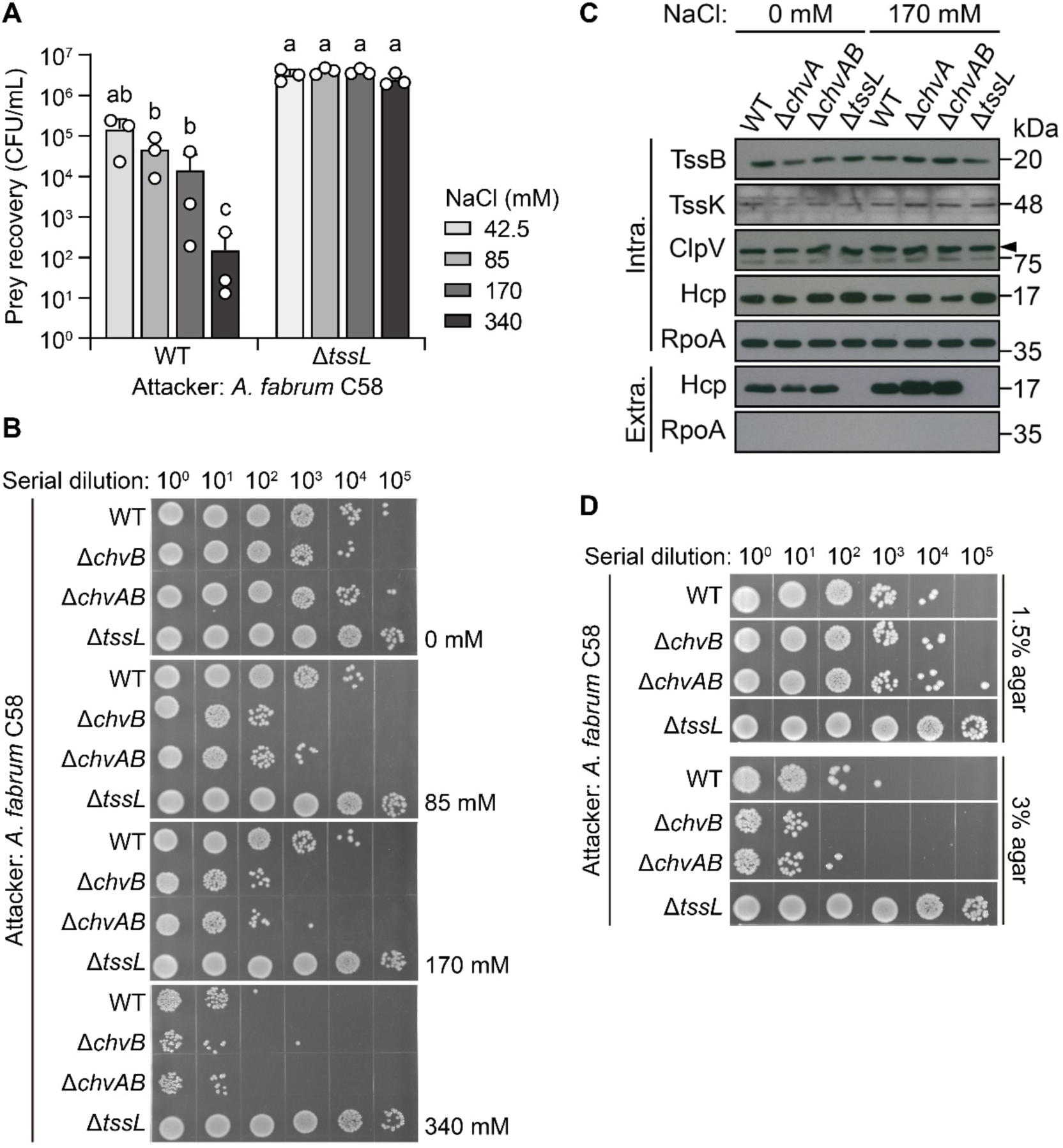
Increased osmolarity and deficiency of cyclic-β-(1,2)-glucan (CβG) promote T6SS antibacterial activity. (**A**) *Agrobacterium* shows stronger interbacterial killing activity upon NaCl supplement. *A. fabrum* C58 wild type (WT) and Δ*tssL* (as a T6SS-negative control) were co-cultured with *E. coli* DH10B prey on 3% LB agar plates containing various NaCl concentrations. The prey cells were recovered, serially diluted, and spotted on a selective medium from biological triplicate, and the calculated colony forming unit (CFU) data were plotted by mean ± SD on a log 10 scale. Statistical significance was indicated on top of the bars by the letter (p<0.05). (**B**) Abolishing CβG production enhances T6SS-dependent killing. *A. fabrum* C58 WT, Δ*tssL*, and two strains deficient in producing CβG (Δ*chvA* and Δ*chvAB*) were mixed with *E. coli* prey as described above. The prey cells were recovered, serially diluted, and spotted on a selective medium. The images were taken after the colony formed. (**C**) Deletion of Δ*chvA* and Δ*chvAB* increases T6SS-dependent secretion. Agrobacterial strains were cultivated in AB-MES with or without an additional 170 mM NaCl. The non-secreted (Intracellular; Intra.) and secreted (Extracellular; Extra.) proteins were collected for western blotting. The immunodetected proteins were labeled on the left with a black wedge if needed. The non-secreted protein RpoA was used as a loading control. (**D**) Surface stiffness and CβG production influence the interbacterial killing activity. Agrobacterial strains were mixed with the prey cells as described above and spotted onto a 1.5% or 3% LB agar plate for the interbacterial killing assay. The images of recovered prey cells were taken as described in Fig. 1B.

Since high osmolarity inhibits the biosynthesis of cyclic-β-(1,2)-glucan (CβG) (24), a periplasmic glucan required for attachment to the plant surface and maintenance of periplasmic osmolarity (25, 26), we then tested whether the increased killing activity is associated with a reduced level of CβG. In *Agrobacterium*, CβG is synthesized in the cytosol by ChvB and is translocated to the periplasm via ChvA (27, 28). Thus, we generated Δ*chvB* single and Δ*chvAB* double mutants and performed an interbacterial killing assay on LB agar plates with various NaCl concentrations (Fig.1B). The results showed that both Δ*chvB* and Δ*chvAB* mutants exhibited higher killing activity compared to WT. The killing activity of both Δ*chvB* and Δ*chvAB* mutants further increased along with the increased NaCl concentrations, suggesting that the increased T6SS-dependent killing is not solely dependent on CβG.

To determine whether the increased killing of Δ*chvB* and Δ*chvAB* mutants is associated with enhanced T6SS activity, we performed a T6SS-dependent secretion assay by growing bacteria in LB media with or without NaCl (Fig.1C). Consistent with the killing assay, more Hcp, a hallmark of T6SS secretion, were detected in the extracellular fractions of Δ*chvB* and Δ*chvAB* in the presence of NaCl. Compared to samples collected from LB with 0 mM NaCl, those with 170 mM NaCl exhibited stronger Hcp signals in the extracellular fractions. Interestingly, the enhanced T6SS-dependent killing phenotype can only be observed on 3%, but not 1.5% LB agar plates (Fig.1D), implying that other factors related to surface stiffness influenced T6SS-mediated competition. These results together suggest that T6SS-dependent secretion and killing are enhanced by hyper-osmolarity and a stiff surface, while the lack of CβG production is not the major contributor to this enhancement.

### Abiotic factors regulate EPS level and T6SS killing

Prior results suggested that glucose enhances T6SS-dependent secretion in *Agrobacterium* (29). However, the *E. coli* survival was much increased when co-cultured with *A. fabrum* C58 WT on the glucose-containing AB-MES plate, suggesting an attenuated T6SS-dependent killing (Fig.2A). We noticed that when grown on glucose-containing agar plates, the agrobacterial colonies became more mucoid, suggesting EPS production. We thus hypothesized that EPS produced by *Agrobacterium* may override the enhanced T6SS-dependent secretion and interfere with its own T6SS-mediated antagonism.

**Fig 2.**
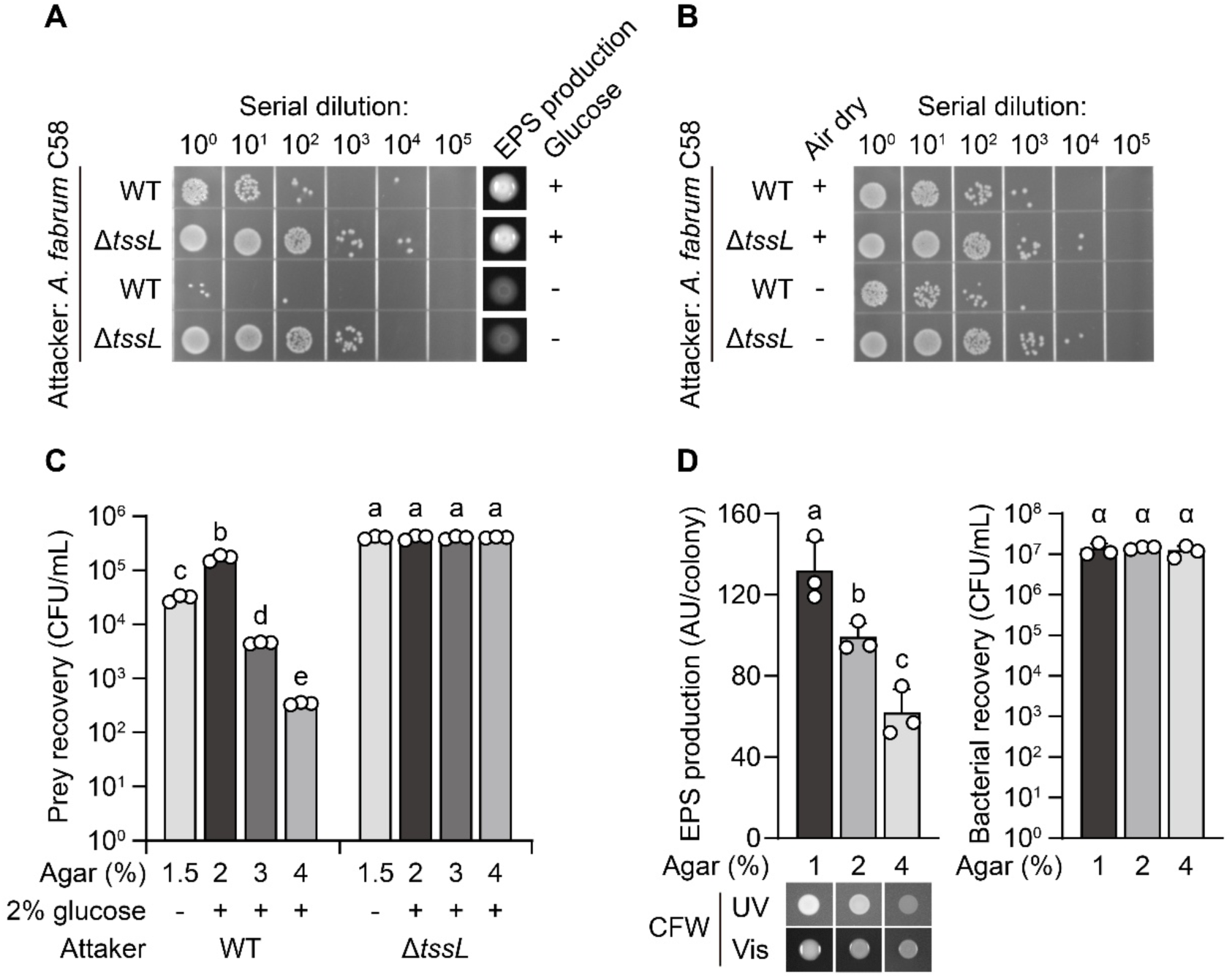
The T6SS-dependent killing and exopolysaccharide (EPS) production are subject to glucose, moisture, and surface stiffness. (**A**) Glucose supplement increases EPS production while decreasing the T6SS-dependent killing in *Agrobacterium*. *A. fabrum* C58 WT and Δ*tssL* were mixed with *E. coli* prey and spotted onto a 3% AB-MES agar plate in the presence (+) or absence (−) of 2% glucose. For the EPS production, agrobacterial cells were spotted on 1.5% ATGN plates with 0.02% Calcofluor white (CFW) stain in the presence (+) or absence (−) of 0.5% glucose. The photos of CFW stained colonies were placed alongside the prey recovery of the interbacterial killing assay, whereas the uncropped images are displayed in Fig. S1A. (**B**) Surface moisture enhances EPS production but reduces the T6SS-dependent killing. The interbacterial killing assay was performed with 1.5% AB-MES agar plates with 2% glucose or 1.5% ATGN plates with 0.02% CFW and 0.5% glucose, respectively. The plates were air-dried (+) for 30 minutes in a laminar flow hood. The images of EPS production assay are displayed in Fig. S1B. (**C**) Increasing agar concentration enhances the T6SS-dependent killing. *A. fabrum* C58 WT and Δ*tssL* were used for an interbacterial killing assay against *E. coli* prey. The bacterial samples were dropped onto AB-MES plates containing different agar concentrations in the presence (+) or absence (−) of 0.5% glucose. The prey recovery from biological triplicate is plotted by mean ± SD on a log 10 scale. The agar concentrations, presence of glucose, and attacker genotypes are indicated below. Statistical significance is indicated on top of the bars by the letter (p<0.05). (**D**) EPS production is decreased by increasing the agar concentration irrelevant of bacterial growth. *A. fabrum* C58 WT was dropped on 0.02% CFW ATGN agar plates containing 1%, 2%, and 4% agar. The CFW signal intensity (arbitrary unit; AU) per colony from the biological triplicate is plotted by mean ± SD. Statistical significance was indicated on top of the bars by the letter (p<0.05). The agrobacterial recovery of each colony was quantified as CFU per mL and plotted alongside the EPS production on a log 10 scale. The photos of the colony under UV or visible light are displayed below the EPS production.

In *Agrobacterium*, the most abundantly secreted EPS is succinoglycan (SG), which can be detected by the fluorescent dye calcofluor white (CFW) (30, 31). We compared CFW fluorescence of WT and Δ*tssL* grown on ATGN plates with or without 0.5% glucose, and found that both strains showed higher fluorescence on the ATGN plate with glucose (Fig.S1A). Besides the glucose depletion, we noticed that the low humidity and a stiff surface led to lower CFW fluorescence, suggesting a reduced SG production (Fig. S1B).

To correlate the EPS production and T6SS-dependent killing, we performed killing assays on AB-MES plates with prolonged plate drying or increased agar concentration (Fig.2B and Fig.2C). As predicted, reduced killing was observed under the conditions in which EPS was highly produced. Such killing outcomes or EPS production were not due to varied bacterial growth, as comparable *A. fabrum* C58 cells were recovered from the ATGN plates containing different agar concentrations (Fig.2D). Taken together, our results indicated that several abiotic factors, including glucose, moisture, and surface stiffness, influence the EPS production and hence agrobacterial T6SS-dependent killing activity.

### SG compromises T6SS killing activity independent of affecting the function of the T6SS machinery

Because SG synthesis in *Agrobacterium* requires *exoA*, and SG production is enhanced in Δ*exoR* (23, 32, 33), we used Δ*exoA* and Δ*exoR* to evaluate the impacts of SG on T6SS-dependent killing (Fig.3A and Fig.3B). We found that Δ*exoA* showed stronger killing activity than WT against *E. coli* or susceptible *A. fabrum* C58, Δ3TIs, which lacks all three toxin-immunity pairs (Ma *et al.*, 2014) (Fig.3A). In contrast, Δ*exoR*-mediated killing is reduced, regardless of whether glucose was supplemented in the culture medium (Fig.3B). Complementation of *exoA* and *exoR* in the respective mutant fully or partially restored the killing phenotype.

**Fig 3.**
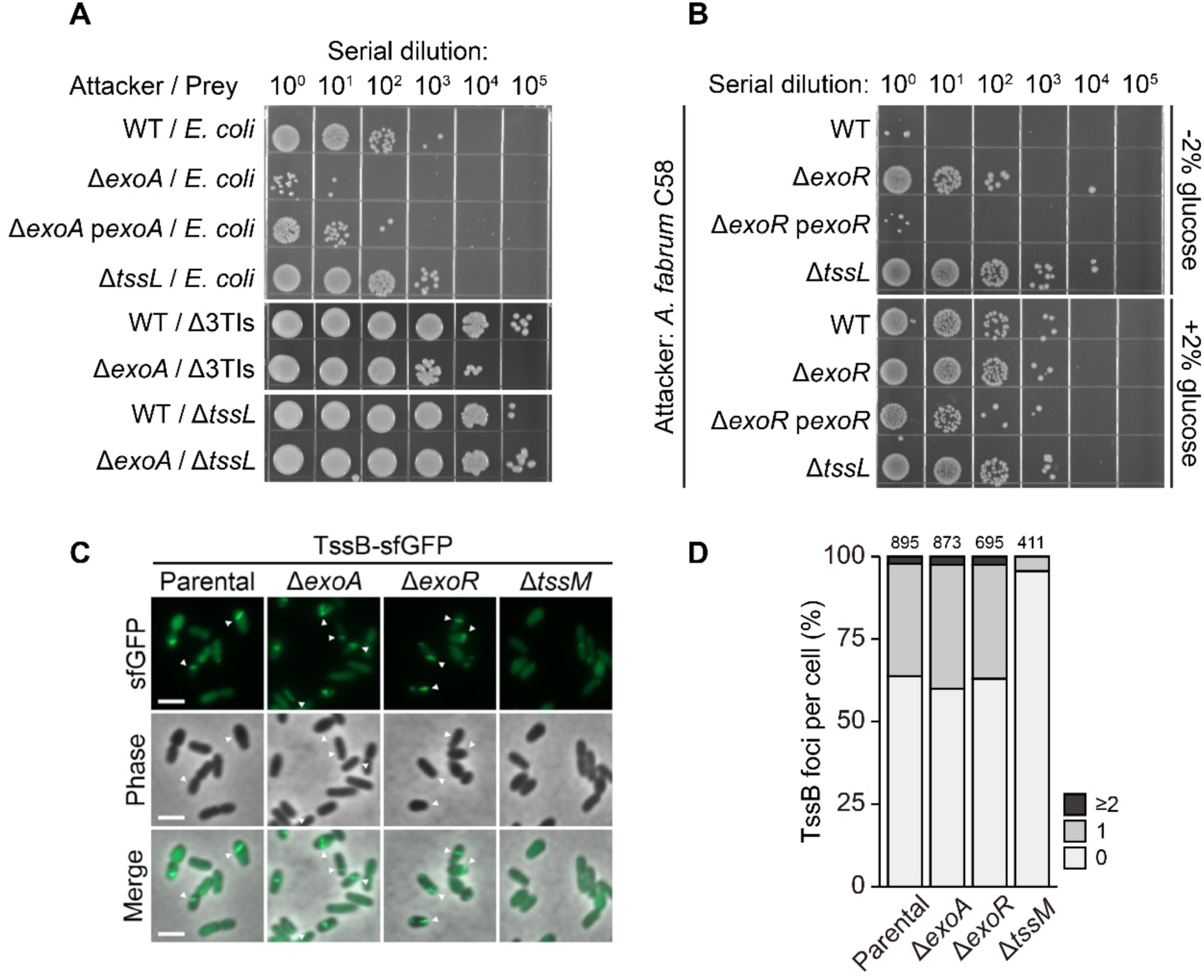
Production of succinoglycan (SG) attenuates T6SS-dependent killing without influencing the T6SS machinery. (**A**) Biosynthesis of SG compromises the self-inflicted T6SS attack during inter- or intra-species competition. *A. fabrum* C58 strains were used as attackers against *E. coli* (top 4 rows), toxin-immunity pairs-deleted *A. fabrum* C58 (Δ3TIs, middle 2 rows), or C58 Δ*tssL* (bottom 2 rows). The images of recovered prey cells were taken as described above. (**B**) Overproduction of SG reduces T6SS-dependent killing regardless of glucose supplement. *A. fabrum* C58 attackers were mixed with *E. coli* prey for the interbacterial killing assay on 2% AB-MES agar plate with or without 2% glucose. The images of prey recovery were displayed beside the genotype of the attackers. For both Fig. 3A and 3B, all attacker strains harbor the pRL662 shuttle vectors, while the strains trans-expressing *exoA* or *exoR* genes were indicated with a prefix “p”. (**C** and **D**) Perturbation of SG biosynthesis does not impact the T6SS apparatus assembly. The formation of the T6SS contractile sheath is shown in Fig. 3C by observing agrobacterial strains expressing TssB-sfGFP with live fluorescent microscopy. The assembled contractile sheaths are indicated by white arrowheads. Scale bar: 2 µm. The quantitative TssB foci formation is shown in Fig. 3D, and the cells used for quantification are indicated at the top.

To determine whether these phenotypic differences reflected altered T6SS activity, we examined Hcp expression of mutant strains. Notably, the level of secreted Hcp from Δ*exoA* and Δ*exoR* was similar to that of WT. Similarly, the levels of non-secreted T6SS proteins were comparable among all strains (Fig.S2). This suggested that the altered killing outcome in Δ*exoA* or Δ*exoR* was independent of T6SS activity.

To assess whether SG production affects T6SS assembly, we observed contractile sheath formation in Δ*exoA* and Δ*exoR* expressing sfGFP-TssB (Fig.3C). As a negative control, we included Δ*tssM,* a strain that lacks a structural component required for apparatus assembly (34). While Δ*tssM* showed no elongated sheath, Δ*exoA* and Δ*exoR* exhibited a comparable percent of TssB sheath structures per cell as WT expressing TssB-sfGFP (Fig.3D). Collectively, we deduced that SG production compromised T6SS-dependent killing without hindering T6SS machinery function.

### SG production overrides T6SS activity in determining the T6SS-killing outcome

Previous work showed that glucose supplementation led to a reduction in T6SS-mediated killing but an increase in T6SS secretion (29). Under these conditions, the reduced killing correlates with increased EPS production, likely due to abundant SG levels (Fig.2A). This led us to predict that Δ*exoA*, the SG synthesis-deficient mutant, will exhibit stronger T6SS-dependent killing in the presence of glucose. As expected, Δ*exoA* conferred a stronger killing as the *E. coli* prey survival against Δ*exoA* was reduced in the glucose-supplemented condition. In contrast, the prey survival against WT showed the opposite trend (Fig.4A). These results support the conclusion that the SG production can override the T6SS activity in determining the overall T6SS-killing outcome.

**Fig 4.**
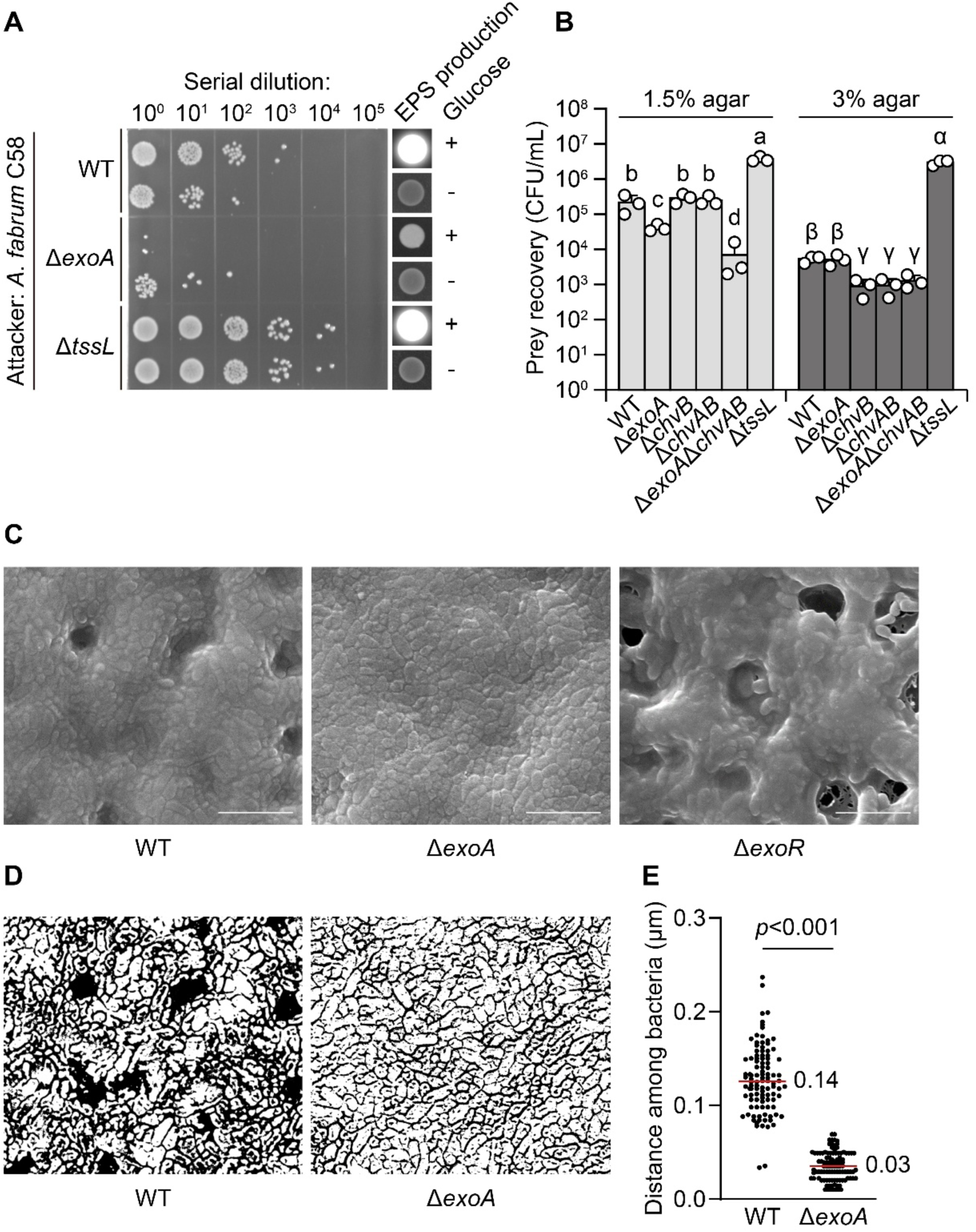
SG production is a major factor governing the interbacterial killing activity. (**A**) The reduction of T6SS-dependent killing by glucose is dependent on SG biosynthesis. The T6SS-killing activity and EPS production of *A. fabrum* C58 WT, Δ*exoA,* and Δ*tssL* were tested as described in Fig. 2A. The uncropped images of EPS production are displayed in Fig. S1C. (**B**) Production of CβG and SG independently reduces T6SS-dependent killing. The interbacterial killing assay using *A. fabrum* strains as attackers against *E. coli* was carried out on 1.5% or 3% LB agar plate. The genotypes of the attackers are indicated in the x-axis, while the prey recovery from three biological triplicate was plotted by mean ± SD in the y-axis on a log 10 scale. Statistical significance was indicated on top of the bars by the letter (p<0.05). (**C**) The scanning electron micrographs of C58 WT, Δ*exoA,* and Δ*exoR*. Scale bar: 5 µm. (**D** and **E**) SG production increases the distance between bacteria. The agrobacterial cells and intercellular spaces of WT and Δ*exoA* from Fig. 4C are displayed with a black-and-white contrast in Fig. 4D. The quantitative distance among bacteria is shown in Fig. 4E. The median is indicated by the red lines and the number alongside.

To dissect the contributions of different EPS species to T6SS killing activity, we generated the single and double mutants of Δ*exoA* and Δ*chvAB* for the interbacterial killing assay (Fig.4B). When the killing assay was carried out on a 3% agar plate, in which SG production was inhibited, Δ*exoA* showed similar antibacterial activity to WT, whereas the Δ*chvB* and Δ*chvAB* mutants exhibited stronger killing. In contrast, when the killing assay was carried out on a 1.5% agar plate, where SG was produced, Δ*exoA*, but not the Δ*chvAB* mutant, showed stronger killing than WT. The antagonism was further enhanced in the Δ*exoA*Δ*chvAB* mutant. Taken together, this result suggests that the production of CβG and SG affects T6SS-dependent killing independently, while SG production is the major factor compromising the T6SS-mediated antagonism.

### SG serves as a physical barrier separating the attackers from the prey

Since T6SS-mediated antagonism requires cell contact, we tested whether the presence of SG is associated with increased cell-to-cell distance, thereby compromising T6SS killing. We used scanning electron microscopy to determine the cell morphology and cell-to-cell distance of WT, Δ*exoA,* and Δ*exoR* (Fig.4C and Fig.4D). We were unable to define the boundaries of Δ*exoR* since the cells were embedded in extracellular fluid. The measurement of distance between cells showed that the WT cells were further away from sister cells, with an average distance of 0.14 μm, compared to the Δ*exoA* cells (0.03 μm on average; Fig.4E). This observation reinforces that SG separates the neighboring cells by increasing the distance between cells.

To test whether the increased distance from the SG production attenuates T6SS killing, we made a mixture of two mutant strains, Δ*tssL*, the T6SS-deficient mutant capable of producing SG, and Δ*exoA* as an attacker population, and paired them against *E. coli* prey (Fig.5A and Fig.5B). We also mixed the Δ*exoA*Δ*tssL* double mutant with Δ*exoA* to generate a population with T6SS-competent cells but no SG-producing cells. As expected, Δ*exoA* mixed with Δ*tssL* exhibited a reduced interbacterial killing compared to that mixed with Δ*exoA*Δ*tssL*. These results suggest that SG produced by the attacker population can indirectly protect neighboring cells, including prey, by increasing cell-to-cell distance and reducing T6SS-mediated killing.

**Fig 5.**
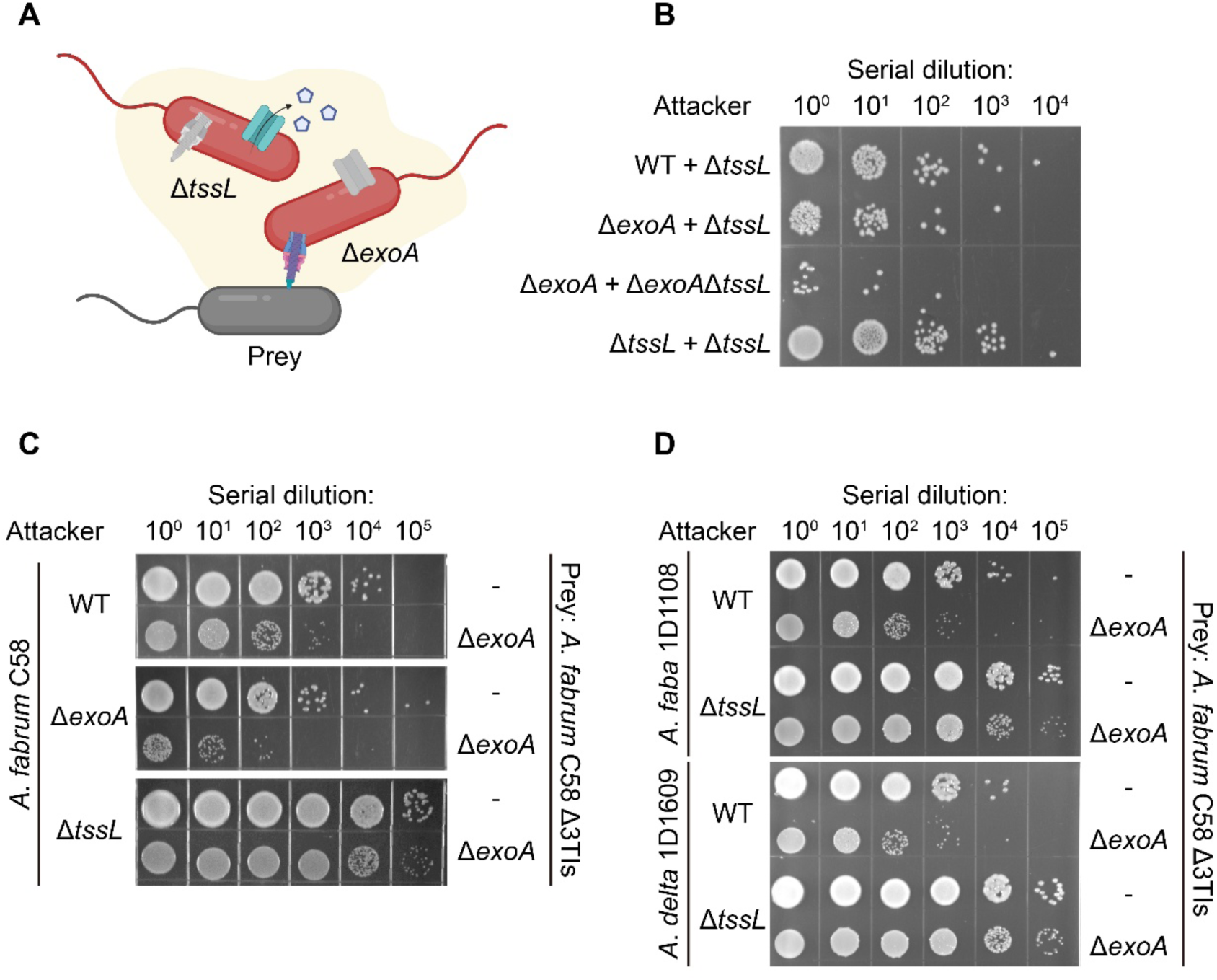
SG confers protection against T6SS killing. (**A**) The schematic of the experimental design for extracellular supplement of SG from Δ*tssL* complements the phenotypes of Δ*exoA* on T6SS-dependent killing. The *A. fabrum* C58 and *E. coli* are colored in red and dark grey, respectively. The inactive T6SS apparatus or SG biosynthesis is in light gray, and the extracellular SG is indicated by pentagons and the yellow layer surrounding the cells. (**B**) Different combinations of agrobacterial strains were used as attackers for the interbacterial killing assay against *E. coli* prey. The image of recovered prey was taken as described above. The attacker strains were mixed at a 1:1 ratio and mixed with E*. coli* to get a final ratio of 20:1 (*A. fabrum* C58: *E. coli*). (**C** and **D**) SG protects *A fabrum* C58 during intra- (**C**) and inter-species (**D**) competition. *A. fabrum* C58, *A. fabacearum* (*A. faba*) 1D1108, and *A. deltaense* (*A. delta*) 1D1609 were used as attackers for the interbacterial competition assay against *A. fabrum* C58 Δ3TIs and its Δ*exoA* derivative. The photos of recovered prey cells are displayed, whereas the genotypes of attackers and prey are indicated on the left or right of the photos, respectively.

Because SG has collateral effects that reduce the efficacy of T6SS-mediated killing within mixed populations, we next asked whether SG produced by *Agrobacterium* cells can protect those cells from T6SS attack by other bacteria. We first performed intraspecies competition using susceptible *A. fabrum* C58 Δ3TIs strain (Ma *et al.*, 2014). We demonstrated that Δ3TIsΔ*exoA* was more susceptible than Δ3TIs to T6SS attack from *A. fabrum* C58 WT, verifying that SG conferred protection against contact-dependent antagonism (Fig.5C). In addition, the survival of Δ3TIs and Δ3TIsΔ*exoA* was lower when competed with Δ*exoA* than against WT, indicating that SG plays a dual role in both defense and offense (Fig.5C). We next tested whether SG protects *A. fabrum* C58 against other agrobacterial strains. Relative to Δ3TIs, the Δ3TIsΔ*exoA* strain was more sensitive to T6SS attack from *A. fabacearum* 1D1108 and *A. deltaense* 1D1609 (Fig.5D). We also investigated the protective effect of SG against *Pseudomonas putida* F1. The survival rates of *A. fabrum* C58 Δ*tssL* were comparable when competing against *P. putida* WT or its Δ*tssM* mutant (Fig.S3A), suggesting that *P. putida* T6SS was not effective at antagonizing C58. However, Δ*exoA*Δ*tssL* was slightly more susceptible to *P. putida* in a T6SS-dependent manner (Fig.S3B). Taken together, we demonstrated that SG protects bacteria from T6SS attacks regardless of attack-prey pairs. This may be due to the physical distancing that impairs the contact-dependent antagonism.

## DISCUSSION

Bacteria coordinate various physiological processes in response to abiotic cues. Using *A. fabrum* C58 as a model, we investigated the interplay among carbon sources, EPS production, and T6SS-mediated competition. We reported that the T6SS-mediated killing is modulated by osmotic stress and polysaccharide production, in which succinoglycan (SG) is the major factor affecting the killing outcome. In combination with current knowledge and our findings, we proposed a model to illustrate our findings (Fig.6). Several environmental cues, including carbon source, osmolarity, and surface stiffness, modulate the production of SG. This secreted EPS confers a slimy peripheral structure that separates *Agrobacterium* from neighboring cells. The physical distancing hence impairs the effectiveness of T6SS-mediated antagonism. In addition to SG, the periplasmic polysaccharide cyclic-β-(1,2)-glucan (CβG) negatively regulates the T6SS-dependent secretion. The production of CβG is subjected to osmolarity and inhibits the expression of SG biosynthesis genes (25). ExoR-ChvG/ChvI signal pathway is a global regulator, which regulates both T6SS and SG in response to acidic signal (21–23). Co-activation of the T6SS antibacterial machinery and SG biosynthesis confers protection to *Agrobacterium* against incoming T6SS attacks, but at the same time compromises the efficacy of its own T6SS-mediated assault. This dual effect likely arises from SG-mediated extracellular matrix formation, which acts as a physical barrier that restricts close cell–cell contact required for effective effector delivery. These findings reveal a tradeoff between EPS-mediated defense and contact-dependent T6SS offense, emphasizing the finely tuned balance between bacterial protection and aggression in the highly competitive polymicrobial rhizosphere.

**Fig 6.**
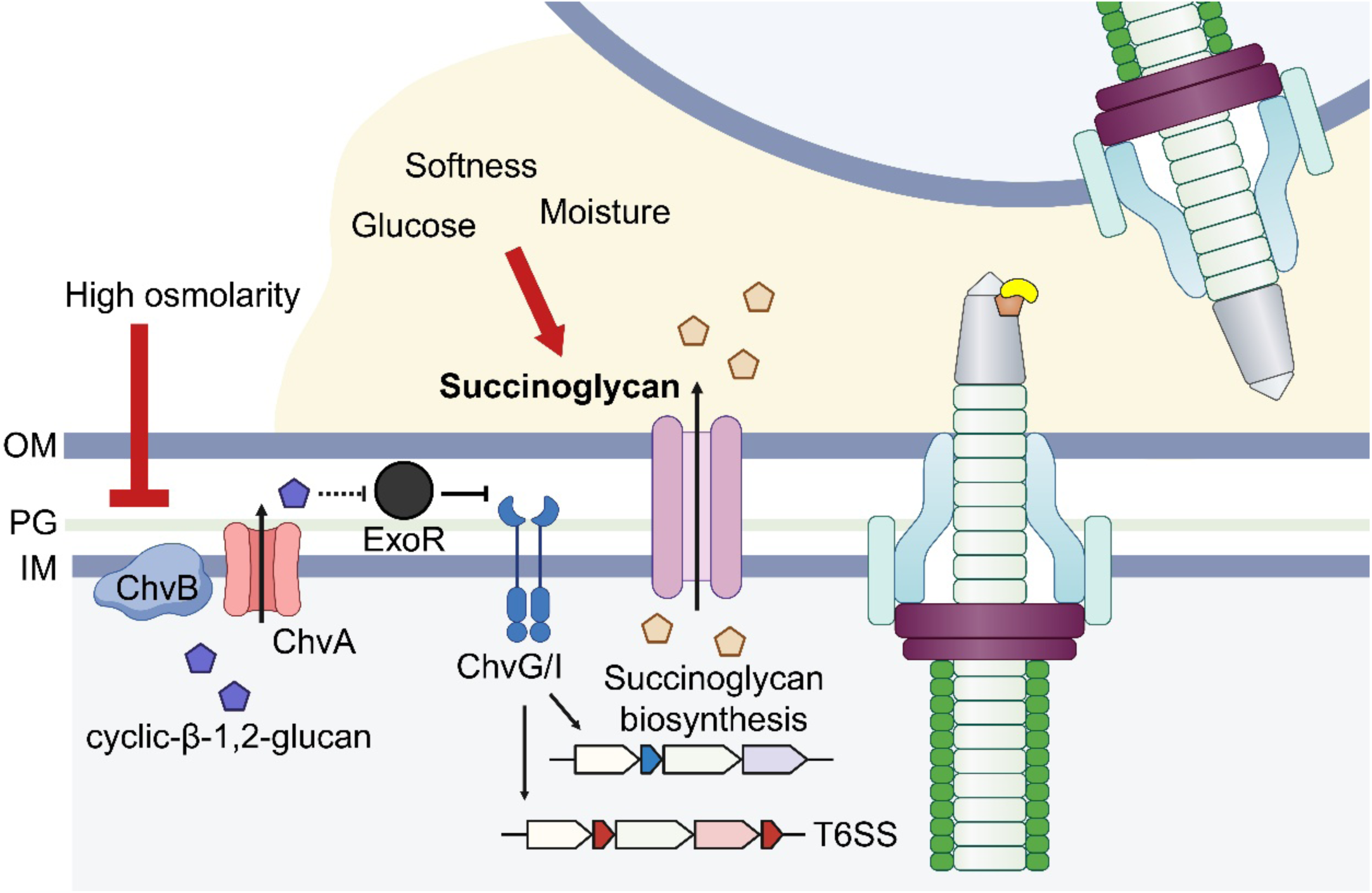
Proposed model for the involvement of cyclic-β-(1,2)-glucan and succinoglycan in T6SS offense and defense in *Agrobacterium*. The T6SS-killing outcome is subject to several environmental factors. Increased osmolarity reduces the cyclic-β-(1,2)-glucan biosynthesis and enhances T6SS-dependent secretion and succinoglycan biosynthesis, through the ExoR-ChvG/ChvI signaling pathway, which also regulates T6SS gene expression. Succinoglycan, in response to glucose supplement, surface stiffness, and moisture, serves as the major factor attenuating the T6SS attack from self and non-self by providing physical distancing. OM: outer membrane. PG: peptidoglycan. IM: inner membrane.

Despite being identified as a common polysaccharide produced by α-Proteobacteria for decades, the physiological roles of SG remain elusive and appear to be diverse among various species. In *Sinorhizobium meliloti*, a nodule-forming symbiont, the succinyl group of SG is required for establishing successful symbiosis (35). However, in *Mesorhizobium loti*, another symbiotic rhizobia, or *Agrobacterium*, SG production is dispensable for nitrogen-fixing nodule formation or crown gall formation, respectively (36, 37). The discrete phenotypes were also observed in the biofilm formation of different SG overproducers (23, 38). The *exoR*::Tn5 mutation in *S. meliloti* leads to increased SG production along with enhanced abiotic biofilm formation (39). In contrast, deletion of *exoR* in *Agrobacterium* confers a complete loss of static biofilm on abiotic surfaces, although this deficiency seemed to result from other genes in the *exoR* regulon (23). These discrepancies suggest that the core function of SG may not be mediating host-microbe interactions or facilitating surface attachment. Interestingly, cell envelope perturbation upregulates the biosynthetic genes of SG in *Agrobacterium* through the ChvG-ChvI signaling pathway (40). Deletion of *chvI* sensitizes *Agrobacterium* to β-lactam antibiotics, suggesting that the ChvG-ChvI signaling pathway mediates stress responses, including SG biosynthesis. Indeed, SG is highly produced in acidic conditions, reflecting the rhizosphere microenvironments, where bacteria face intense competition and selective pressure from the host. As a result, extracellular matrices are potentially produced for defense and survival.

In this study, we demonstrated that SG production protects *Agrobacterium* against external T6SS attack. However, such production also compromises the T6SS attack originating from itself. As *Agrobacterium* coordinates the expression of the SG biosynthetic and T6SS-encoding genes, we questioned the ecological implications. Although T6SS is often framed as an offensive weapon when viewed at the single-cell level, studies also suggest its defensive contribution to niche establishment and maintenance (41, 42). After initial colonization and clonal expansion, T6SS can eliminate invaders and maintain the integrity within a colony (43). Moreover, T6SS has been shown to “punish” cheaters within microbial communities. For example, the *Yersinia* T6SS-1 effector TepC exhibits versatile functions as a siderophore and DNase to prevent resource foraging by non-kin cells (44). As the secreted EPS is a public good that provides collective protection and resilience (12, 40), T6SS-dependent killing may function similarly to restrict the growth of non-kin cells not producing EPS. Overall, our findings uncover a sophisticated balance in the regulation of bacterial competition, positioning SG in the offense-defense dynamics of *Agrobacterium*. The coordinated expression of T6SS and SG supports a model in which T6SS functions not only as an offensive weapon but also as a defensive and community-stabilizing mechanism in environments.

## MATERIALS AND METHODS

### Bacterial strains, plasmids, and growth conditions

All bacterial strains and plasmids used in this study are listed in Table S1. If not specifically indicated, the agrobacterial strains were grown in 523 medium at 25°C (45), while *Pseudomonas putida* and *Escherichia coli* strains were grown in LB medium at 30 °C and 37 °C, respectively. Antibiotics were added if needed with the following concentrations: gentamycin, 15 μg/mL for *P. putida* and 50 μg/mL for *Agrobacterium* spp.; irgasan (triclosan), 25 ug/mL for *P. putida*; spectinomycin, 100 μg/mL for *E. coli*.

### Construction of plasmids and mutants

For the trans complementation, the gene of interest was amplified by PCR, followed by restriction digestion and ligation into the pRL662 shuttle vector. For the in-frame deletion, the double crossover method was used via the pJQ200ks suicide plasmid as described previously (46). To generate *retS* and *tssM* deletion mutants in *P. putida*, mutant constructs cloned into pEXG2 were transformed into *E. coli* S17-1 (47). *E. coli* S17-1 carrying the mutant constructs and the recipient *P. putida* strains were grown overnight at 30 °C on LB plates containing appropriate antibiotics. Cells were then scraped from the plates and mixed at a 10:1 donor-to-recipient ratio for each mating pair. The mixtures were spread onto LB agar plates and incubated at 30 °C for 6 h to facilitate plasmid transfer by conjugation. The cell mixtures were then scraped into LB medium and plated on LB agar containing irgasan (triclosan) and gentamicin to select for *P. putida* cells with integrated plasmids. The resulting merodiploid *P. putida* strains were grown overnight in non-selective LB medium at 30 °C, followed by counter-selection on LB agar supplemented with 5% (*w/v*) sucrose. Gentamicin-sensitive, sucrose-resistant colonies were screened for allelic replacement by colony PCR, and the mutations were confirmed by Sanger sequencing of the PCR products.

### Secretion assay and interbacterial killing assay

The secretion assay was executed following the protocol from our previous publication (46) using AB-MES medium (pH 5.5) (18). For the interbacterial killing assay, the attacker and prey cells were collected and mixed as described previously (46). The bacterial mixtures were then spotted onto LB or AB-MES agar plates with various concentrations of agar, NaCl, or glucose for experimental purposes.

### Calcofluor white stain

The Calcofluor white stain was carried out as described (23) with minor modifications. *Agrobacterium* cells grown in 523 medium overnight were harvested and adjusted to an OD_600_ of 2 in 0.9% saline. Two μL of suspension was spotted onto ATGN agar plates containing 0.02 % calcofluor white (Sigma-Aldrich, USA) and incubated at 25°C for 3 to 4 days. The colonies were imaged by ultraviolet or visible light with the Gel Doc XR+ Gel Documentation System (Bio-Rad, USA). The signal intensity was determined by the ImageJ package (48).

### Cryo Scanning Electron Microscopy (cryo-SEM)

Overnight cultured agrobacterial cells were centrifuged and subcultured at an OD_600_ of 0.2 in AB-MES for 1 day. Bacterial cells were harvested and spotted onto an AB-MES agar plate for further growth of 16 to 18 hours. Afterwards, the agrobacterial cells were collected, mounted onto a copper holder, and plunged into liquid nitrogen. The specimens were transferred onto the cold stage, maintained at –95°C, and imaged using SEM (FEI Quanta 200/Quorum PP2000TR FEI).

### Fluorescent microscopy

The *exoA* or *exoR* deletion was introduced in *A. fabrum* C58 expressing C-terminal superfolder green fluorescent protein (sfGFP)-tagged TssB. Fluorescent microscopy was executed as described previously (49), and foci number quantification was performed by MicrobeJ (50, 51).

### Statistical analyses

For the interbacterial killing assay, the prey recovery was subjected to log10 transformation for statistical analysis. For multiple comparisons, samples were analyzed by ANOVA followed by Tukey’s HSD test. The significance was designated when the p-value was lower than 0.05. The two-tailed Student-t test was used for comparisons of two samples. All the statistical analyses were in the Prism 10.4.1 software with the default parameters.

## ACKNOWLEDGEMENT

The authors thank the former and current Lai lab members for their discussion and suggestions. We also acknowledge Dr. Wann-Neng Jen at the Electron Microscope Division of Cell Biology Core for performing cryo-SEM, Ji-Ying Huang for technical assistance on ImageJ analysis, and Genomic Technology Core for Sanger sequencing, all located in the Institute of Plant and Microbial Biology, Academia Sinica. The research conducted in the Lai lab has been supported by an intramural grant from the Institute of Plant and Microbial Biology, Academia Sinica. J.H.C is supported by a grant from the Plant-Biotic Interactions program, project award number 2022-67013-36883 from the U.S. Department of Agriculture’s National Institutes of Food and Agriculture. Any opinions, findings, conclusions, or recommendations expressed in this publication are those of the authors and should not be construed to represent any official USDA or U.S. Government determination or policy.

## AUTHOR CONTRIBUTIONS

YCW and EML contributed to the conceptualization of the project. YCW and LKS contributed to the investigation, methodology, data curation, formal analysis, and data visualization. SYT and CFW contributed to the reagents and methodology. EML and JHC supervised the research. EML managed the project and acquired funding. EML and LKS wrote the initial manuscript draft, with YCW contributing the methodology and figures. Generative AI was used solely for proofreading. All authors reviewed and edited the manuscript.

---

**Fig S1.**
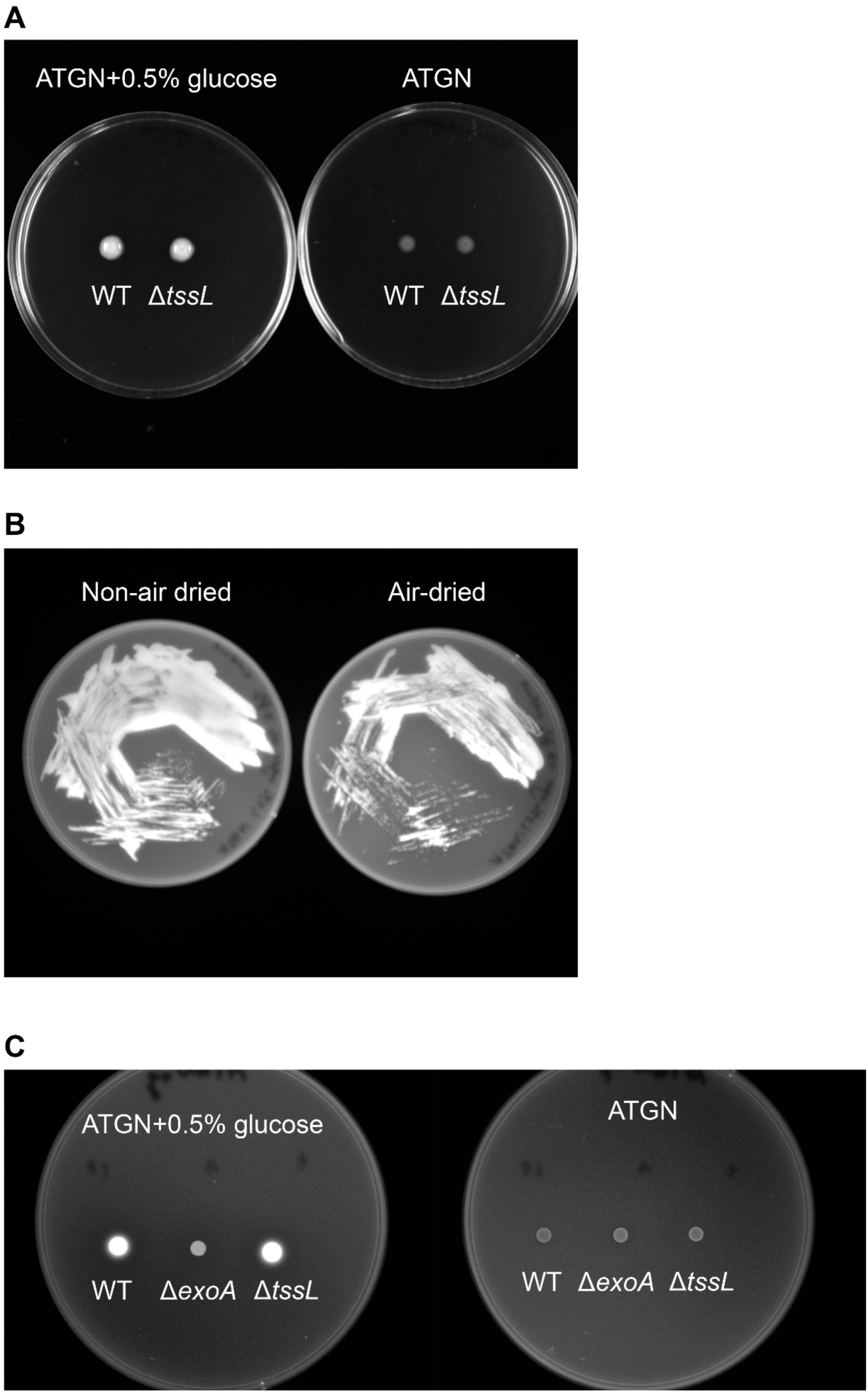
The uncropped images of EPS production assay, related to Fig. 2A (**A**), Fig. 2B (**B**), and Fig. 4A (**C**), respectively. For Fig. S1A and S1C, the used agrobacterial genotypes are labeled below, whereas in Fig. S1B, *A. fabrum* C58 WT was used. The conditions of each experimental set are indicated on top.

**Fig S2.**
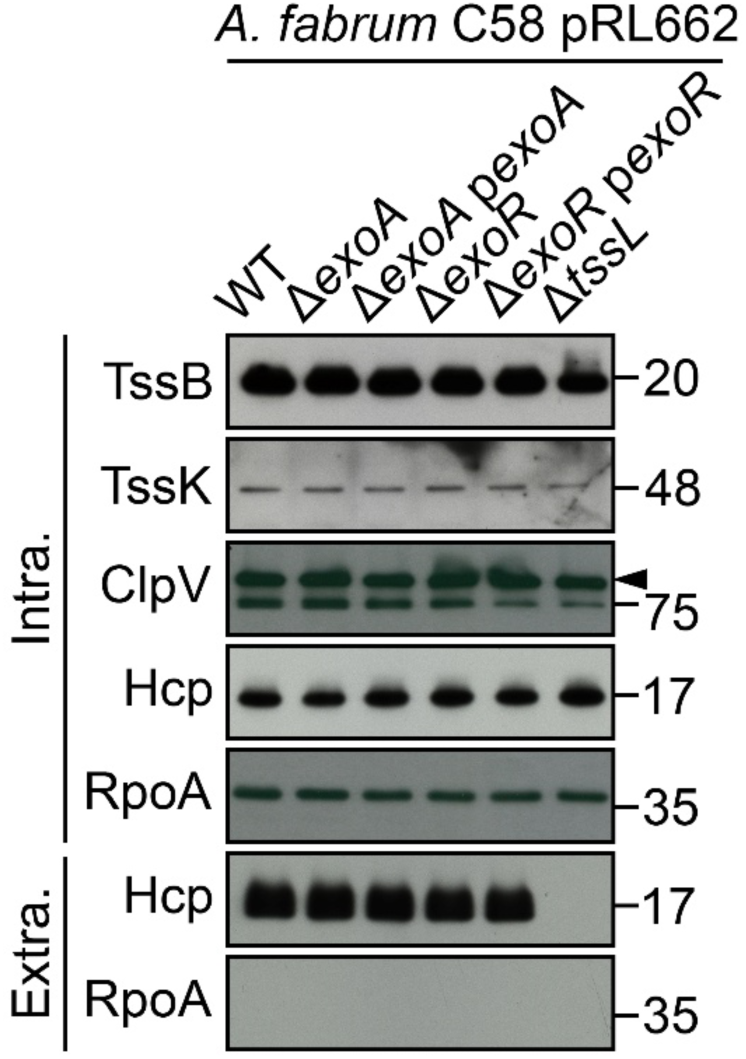
Disruption of SG biosynthesis does not abolish T6SS-dependent secretion. Agrobacterial strains were cultivated in AB-MES for the T6SS-dependent secretion assay. The non-secreted (Cellular) and secreted (Extracellular) proteins were collected for western blotting. The immunodetected proteins were labeled on the left with a black wedge if needed. The non-secreted protein RpoA was used as a loading control.

**Fig S3.**
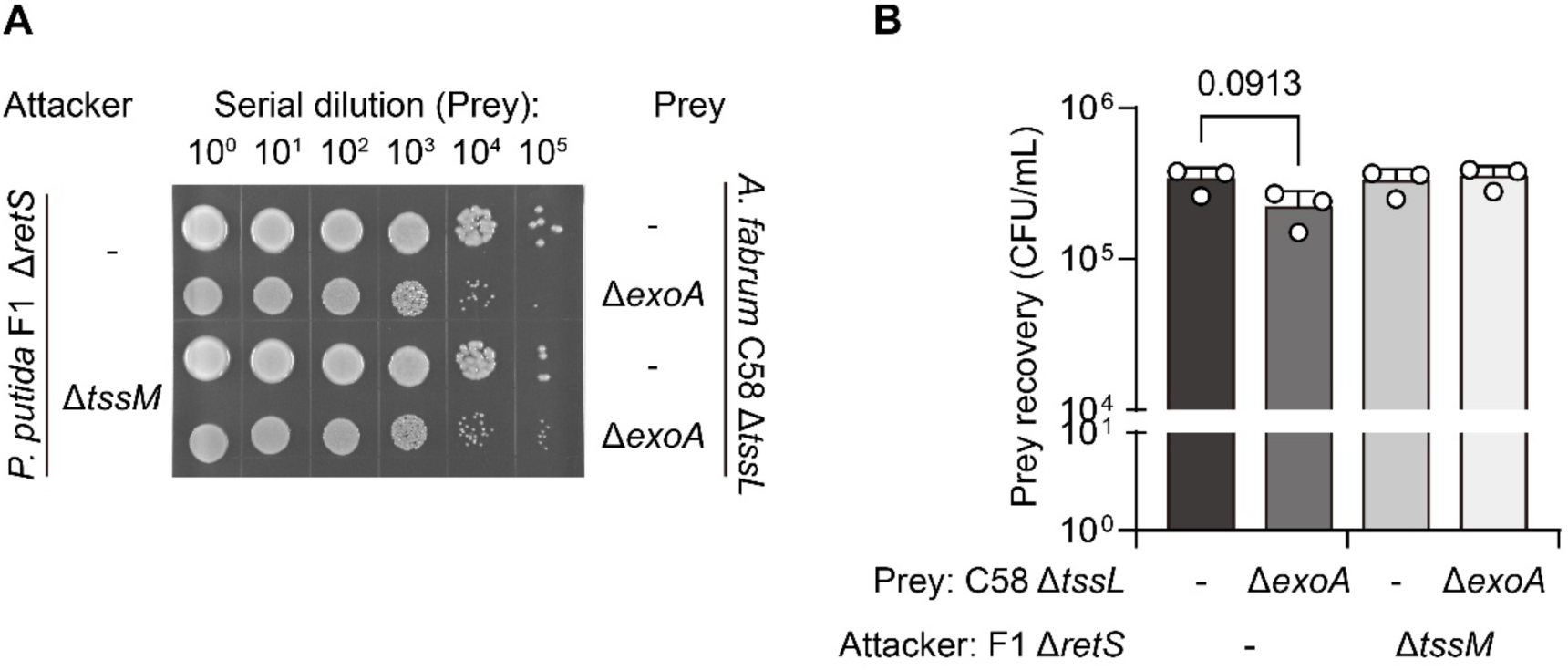
SG biosynthesis confers protection during interspecies competition. (**A**) *A. fabrum* C58 was used as the prey cell against *Pseudomonas putida* KT2440 in the interbacterial competition assay. The photos of recovered prey cells are displayed, while the genotypes of the attackers and prey are indicated on the left and right of the photos, respectively. The quantitative prey recovery from the biological triplicate was plotted by mean ± SD on a log 10 scale and is shown in (**B**). The genotypes of attackers and prey cells are displayed below.

**TABLE S1.** Bacterial strains and plasmids.

| Strain or plasmid | Relevant characteristics | Source or Reference | Lab strain name |
| --- | --- | --- | --- |
| *Agrobacterium* strains |  |  |  |
| *A. fabrum* C58 | Wild-type virulent strain containing nopaline-type Ti plasmid pTiC58 | Eugene Nester | EML530 |
| *A. deltaense* 1D1609 | Wild-type virulent strain containing octopine-type Ti plasmid pTi1D1609 | [1] | EML304 |
| *A. fabacearum* 1D1108 | Wild type virulent nopaline type strain isolated from a Euonymus gall | Clarence I Kado | EML302 |
| C58 $\Delta chvB$ | *chvB* in-frame deletion mutant | This study | EML5724 |
| C58 $\Delta chvAB$ | deletion of *chvAB* genes | This study | EML5705 |
| C58 $\Delta tssL$ | *tssL* in-frame deletion mutant | [2] | EML1073 |
| C58 $\Delta exoA$ | *exoA* in-frame deletion mutant | This study | EML5704 |
| C58 $\Delta exoR$ | *exoR* in-frame deletion mutant | [3] | EML1654 |
| C58 $\Delta 3TIs$ | in-frame deletion of all three toxin-immunity pairs | [2] | EML3561 |
| C58 $\Delta exoA \Delta chvAB$ | *exoA chvAB* in-frame deletion mutant | This study | EML5707 |
| C58 $\Delta 3TIs \Delta exoA$ | *exoA* in-frame deletion in $\Delta 3TIs$ mutant | This study | EML5688 |
| C58 $\Delta tssL \Delta exoA$ | *exoA* in-frame deletion in $\Delta tssL$ mutant | This study | EML5689 |
| *E. coli* strains |  |  |  |
| DH10B | Host for DNA cloning, killing assay | Invitrogen | EML455 |
| S17-1 | Used for plasmid conjugation | Ting, See-Yeun | EML124 |
| *Pseudomonas putida* F1 |  |  |  |
| $\Delta retS$ | *retS* in-frame deletion mutant | Ting, See-Yeun | EML5729 |
| $\Delta retS \Delta tssM$ | *retS tssM* in-frame deletion mutant | Ting, See-Yeun | EML5730 |
| Plasmids |  |  |  |
| pRL662 | GmR, broad-host-range vector derived from pBBR1MCS-2 | [4] | EML315 |
| pTrc200 | SmR SpR, pSV1 origin lacIq, trc promoter expression vector | [5] | EML904 |
| pJQ200KS | GmR, suicide plasmid containing GmR and *sacB* gene for selection of double crossover | [6] | EML121 |
| pRL-*exoA* | GmR, pRL662 expressing *exoA* driven by *lacZp* | This study | EML5726 |
| pRL-*exoR* | GmR, pRL662 expressing *exoR* driven by *lacZp* | [3] | EML1617 |
| pJQ200KS-*exoA* | GmR, used in generating *exoA* deletion mutant of C58 | This study | EML5699 |
| pJQ200KS-*chvB* | GmR, used in generating entire *chvB* deletion mutant of C58 | This study | EML5725 |
| pJQ200KS-*chvAB* | GmR, used in generating entire *chvAB* deletion mutant of C58 | This study | EML5711 |
| pEXG2_PPF1_Δ*retS* | GmR, used in generating entire *retS* deletion mutant of F1 | This study |  |
| pEXG2_PPF1_Δ*tssM* | GmR, used in generating entire *tssM* deletion mutant of F1 | This study |  |

## REFERENCES

1. Palmer JD, Foster KR. Bacterial species rarely work together. Science. 2022;376(6593):581–2.

2. Cianfanelli FR, Monlezun L, Coulthurst SJ. Aim, load, fire: the type VI secretion system, a bacterial nanoweapon. Trends Microbiol. 2016;24(1):51–62.

3. Scouten JM, Hsieh S-C, Sung L-K, Wen Y-HV, Kuo C-H, Lai E-M, et al. Function, Evolution, and Ecology of Type VI Secretion Systems of Plant-Associated Bacteria. Annu Rev Phytopathol. 2025;63.

4. Hespanhol JT, Nobrega-Silva L, Bayer-Santos E. Regulation of type VI secretion systems at the transcriptional, posttranscriptional and posttranslational level. Microbiology (Reading). 2023;169(8).

5. Speare L, Woo M, Bultman KM, Mandel MJ, Wollenberg MS, Septer AN. Host-like conditions are required for T6SS-mediated competition among *Vibrio fischeri* light organ symbionts. mSphere. 2021;6(4):10.1128/msphere.01288-20.

6. Bernardy EE, Turnsek MA, Wilson SK, Tarr CL, Hammer BK. Diversity of clinical and environmental isolates of Vibrio cholerae in natural transformation and contact-dependent bacterial killing indicative of type VI secretion system activity. Applied and environmental microbiology. 2016;82(9):2833–42.

7. Wu CF, Weisberg AJ, Davis EW, 2nd, Chou L, Khan S, Lai EM, et al. Diversification of the Type VI Secretion System in Agrobacteria. mBio. 2021;12(5):e0192721.

8. Coulthurst S. The Type VI secretion system: a versatile bacterial weapon. Microbiology (Reading). 2019;165(5):503–15.

9. Martinkus J, Ginibre-Supersac N, Rivard N, Belot L, Brillard J, Cascales E, et al. Poly-immunity arrays associated with Rhs toxins confer wide protection against competitors. Current Biology. 2026;36(1):176–87. e3.

10. Azhieh A, Hernandez P, Anderson AC, Sychantha D, Verster AJ, Whitney JC, et al. Rapidly evolving orphan immunity genes protect human gut bacteria from intoxication by the type VI secretion system. bioRxiv. 2025.

11. Toska J, Ho BT, Mekalanos JJ. Exopolysaccharide protects Vibrio cholerae from exogenous attacks by the type 6 secretion system. Proc Natl Acad Sci U S A. 2018;115(31):7997–8002.

12. Granato ET, Smith WP, Foster KR. Collective protection against the type VI secretion system in bacteria. ISME J. 2023;17(7):1052–62.

13. Hersch SJ, Watanabe N, Stietz MS, Manera K, Kamal F, Burkinshaw B, et al. Envelope stress responses defend against type six secretion system attacks independently of immunity proteins. Nature microbiology. 2020;5(5):706–14.

14. Tsai C-E, Wang F-Q, Yang C-W, Yang L-L, Nguyen TV, Chen Y-C, et al. Surface-mediated bacteriophage defense incurs fitness tradeoffs for interbacterial antagonism. EMBO J. 2025;44(9):2473.

15. Flaugnatti N, Bader L, Croisier-Coeytaux M, Blokesch M. Capsular polysaccharide restrains type VI secretion in *Acinetobacter baumannii*. eLife. 2025;14:e101032.

16. Hwang HH, Yu M, Lai EM. Agrobacterium-Mediated Plant Transformation: Biology and Applications: American Society of Plant Biologists; 2017.

17. Weisberg AJ, Wu Y, Chang JH, Lai EM, Kuo CH. Virulence and Ecology of Agrobacteria in the Context of Evolutionary Genomics. Annu Rev Phytopathol. 2023;61:1–23.

18. Ma LS, Hachani A, Lin JS, Filloux A, Lai EM. Agrobacterium tumefaciens deploys a superfamily of type VI secretion DNase effectors as weapons for interbacterial competition in planta. Cell Host Microbe. 2014;16(1):94–104.

19. Wu CF, Santos MNM, Cho ST, Chang HH, Tsai YM, Smith DA, et al. Plant-Pathogenic Agrobacterium tumefaciens Strains Have Diverse Type VI Effector-Immunity Pairs and Vary in In-Planta Competitiveness. Molecular Plant-Microbe Interactions. 2019;32(8):961–71.

20. Chou L, Lin YC, Haryono M, Santos MNM, Cho ST, Weisberg AJ, et al. Modular evolution of secretion systems and virulence plasmids in a bacterial species complex. BMC Biol. 2022;20(1):16.

21. Wu CF, Lin JS, Shaw GC, Lai EM. Acid-induced type VI secretion system is regulated by ExoR-ChvG/ChvI signaling cascade in Agrobacterium tumefaciens. PLoS Pathog. 2012;8(9):e1002938.

22. Greenwich JL, Heckel BC, Alakavuklar MA, Fuqua C. The ChvG-ChvI Regulatory Network: A Conserved Global Regulatory Circuit Among the Alphaproteobacteria with Pervasive Impacts on Host Interactions and Diverse Cellular Processes. Annu Rev Microbiol. 2023;77:131–48.

23. Tomlinson AD, Ramey-Hartung B, Day TW, Merritt PM, Fuqua C. Agrobacterium tumefaciens ExoR represses succinoglycan biosynthesis and is required for biofilm formation and motility. Microbiology (Reading). 2010;156(Pt 9):2670–81.

24. Miller KJ, Kennedy EP, Reinhold VN. Osmotic adaptation by gram-negative bacteria: possible role for periplasmic oligosaccharides. Science. 1986;231(4733):48–51.

25. Matthysse AG. Attachment of Agrobacterium to plant surfaces. Front Plant Sci. 2014;5:252.

26. Cangelosi GA, Martinetti G, Nester EW. Osmosensitivity phenotypes of Agrobacterium tumefaciens mutants that lack periplasmic beta-1,2-glucan. J Bacteriol. 1990;172(4):2172–4.

27. Cangelosi GA, Martinetti G, Leigh JA, Lee CC, Thienes C, Nester EW. Role for [corrected] Agrobacterium tumefaciens ChvA protein in export of beta-1,2-glucan. J Bacteriol. 1989;171(3):1609–15.

28. Guidolin LS, Arce-Gorvel V, Ciocchini AE, Comerci DJ, Gorvel JP. Cyclic beta-glucans at the bacteria-host cells interphase: One sugar ring to rule them all. Cell Microbiol. 2018;20(6):e12850.

29. Yu M, Wang Y-C, Huang C-J, Ma L-S, Lai E-M, Comstock LE. Agrobacterium tumefaciens Deploys a Versatile Antibacterial Strategy To Increase Its Competitiveness. J Bacteriol. 2021;203(3):e00490–20.

30. Matthysse AG. Exopolysaccharides of Agrobacterium tumefaciens. Curr Top Microbiol Immunol. 2018;418:111–41.

31. Thompson MA, Onyeziri MC, Fuqua C. Function and Regulation of Agrobacterium tumefaciens Cell Surface Structures that Promote Attachment. Curr Top Microbiol Immunol. 2018;418:143–84.

32. Cangelosi GA, Hung L, Puvanesarajah V, Stacey G, Ozga DA, Leigh JA, et al. Common loci for Agrobacterium tumefaciens and Rhizobium meliloti exopolysaccharide synthesis and their roles in plant interactions. J Bacteriol. 1987;169(5):2086–91.

33. Wu D, Li A, Ma F, Yang J, Xie Y. Genetic control and regulatory mechanisms of succinoglycan and curdlan biosynthesis in genus Agrobacterium. Appl Microbiol Biotechnol. 2016;100(14):6183–92.

34. Durand E, Nguyen VS, Zoued A, Logger L, Pehau-Arnaudet G, Aschtgen M-S, et al. Biogenesis and structure of a type VI secretion membrane core complex. Nature. 2015;523(7562):555–60.

35. Cheng H-P, Walker GC. Succinoglycan is required for initiation and elongation of infection threads during nodulation of alfalfa by *Rhizobium meliloti*. J Bacteriol. 1998;180(19):5183–91.

36. Cangelosi G, Hung L, Puvanesarajah V, Stacey G, Ozga D, Leigh J, et al. Common loci for *Agrobacterium tumefaciens* and *Rhizobium meliloti* exopolysaccharide synthesis and their roles in plant interactions. J Bacteriol. 1987;169(5):2086–91.

37. Kelly SJ, Muszyński A, Kawaharada Y, Hubber AM, Sullivan JT, Sandal N, et al. Conditional requirement for exopolysaccharide in the *Mesorhizobium*–Lotus symbiosis. Molecular plant-microbe interactions. 2013;26(3):319–29.

38. Fujishige NA, Kapadia NN, De Hoff PL, Hirsch AM. Investigations of *Rhizobium* biofilm formation. FEMS microbiology ecology. 2006;56(2):195–206.

39. Wells DH, Chen EJ, Fisher RF, Long SR. ExoR is genetically coupled to the ExoS–ChvI two-component system and located in the periplasm of *Sinorhizobium meliloti*. Molecular microbiology. 2007;64(3):647–64.

40. Williams MA, Bouchier JM, Mason AK, Brown PJ. Activation of ChvG-ChvI regulon by cell wall stress confers resistance to β-lactam antibiotics and initiates surface spreading in *Agrobacterium tumefaciens*. PLoS Genet. 2022;18(12):e1010274.

41. Stubbusch AK, Peaudecerf FJ, Lee KS, Paoli L, Schwartzman J, Stocker R, et al. Antagonism as a foraging strategy in microbial communities. Science. 2025;388(6752):1214–7.

42. Gallegos-Monterrosa R, Coulthurst SJ. The ecological impact of a bacterial weapon: microbial interactions and the Type VI secretion system. FEMS MICROBIOL REV. 2021;45(6).

43. Hersch SJ, Manera K, Dong TG. Defending against the type six secretion system: beyond immunity genes. Cell Rep. 2020;33(2).

44. Song L, Xu L, Wu T, Shi Z, Kareem HA, Wang Z, et al. Trojan horselike T6SS effector TepC mediates both interference competition and exploitative competition. ISME J. 2024;18(1):wrad028.

45. Kado CI, and Heskett, M.G.. Selective media for isolation of *Agrobacterium, Carynebacterium, Erwinia, Pseudomonas*, and *Xanthomonas*. . Phytopathology. 1970;60:969–76.

46. Lin JS, Ma LS, Lai EM. Systematic Dissection of the Agrobacterium Type VI Secretion System Reveals Machinery and Secreted Components for Subcomplex Formation. PLoS One. 2013;8(7):e67647.

47. Rietsch A, Vallet-Gely I, Dove SL, Mekalanos JJ. ExsE, a secreted regulator of type III secretion genes in *Pseudomonas aeruginosa*. Proc Natl Acad Sci U S A. 2005;102(22):8006–11.

48. Schindelin J, Arganda-Carreras I, Frise E, Kaynig V, Longair M, Pietzsch T, et al. Fiji: an open-source platform for biological-image analysis. Nat Methods. 2012;9(7):676–82.

49. Wu C-F, Lien Y-W, Bondage D, Lin J-S, Pilhofer M, Shih Y-L, et al. Effector loading onto the VgrG carrier activates type VI secretion system assembly. EMBO Rep. 2020;21(1):e47961.

50. Ducret A, Quardokus EM, Brun YV. MicrobeJ, a tool for high throughput bacterial cell detection and quantitative analysis. Nature microbiology. 2016;1(7):1–7.

51. Bernal P, Furniss RCD, Fecht S, Leung RC, Spiga L, Mavridou DA, et al. A novel stabilization mechanism for the type VI secretion system sheath. Proc Natl Acad Sci U S A. 2021;118(7):e2008500118.

